# Direct sulphate-TOR signalling controls transcriptional reprogramming for shoot apex activation in *Arabidopsis*

**DOI:** 10.1101/2021.03.09.434511

**Authors:** Yongdong Yu, Zhaochen Zhong, Liuyin Ma, Chengbin Xiang, Ping Xu, Yan Xiong

**Affiliations:** Shanghai Centre for Plant Stress Biology, Chinese Academy of Sciences Centre for Excellence in Molecular Plant Sciences, Chinese Academy of Sciences, Shanghai 201602, China; Basic Forestry and Proteomics Research Center, Fujian Agriculture and Forestry University, Fuzhou, Fujian Province 350002, China; University of Chinese Academy of Sciences, 100049 Beijing, China; School of Life Sciences, University of Science and Technology of China, Hefei, Anhui Province 230027, China

## Abstract

Photosynthetic plants play a primary role for the global sulphur cycle in the earth ecosystems by reduction of inorganic sulphate from the soil to organic sulphur-containing compounds. How plants sense and transduce the sulphate availability in soil to mediate their growth remains largely unclear. The target of rapamycin (TOR) kinase is an evolutionarily conserved master regulator of nutrient sensing and metabolic signalling to control cell proliferation and growth in all eukaryotes. Here, we discovered that inorganic sulphate exhibits higher potency than organic cysteine and glutathione for activation of TOR and cell proliferation in the leaf primordium to promote true leaf development in *Arabidopsis*. Chemical genetic analyses further revealed that this sulphate activation of TOR is independent of the sulphate-assimilation process and glucose-energy signalling. Significantly, tissue specific transcriptome analyses uncovered previously unknown sulphate-orchestrating genes involved in DNA replication, cell proliferation, autophagy and various secondary metabolism pathways, which are completely depending on TOR signalling. Systematic comparison between the sulphate- and glucose-TOR controlled transcriptome further revealed that, as the central growth integrator, TOR kinase can sense different upstream nutrient signals to control both shared and unique transcriptome networks, therefore, precisely modulate plant proliferation, growth and stress responses.

## Introduction

Sulphur is an essential nutrient for plants that could be incorporated into sulphur-containing amino acids, glutathione and various secondary metabolites to sustain growth, development and stress adaptation^1-3^. Sulphur availability in soil, mainly in the form of sulphate, is one of major limiting factors for plant growth and crop production. Understanding how plants sense, assimilate and transport soil-absorbed sulphate is essential for precisely modern crop modification, especially provoked by a current reduction in atmospheric sulphur deposition and the lack of available sulphur fertilizers. The functions of transport proteins and enzymes involved in sulphate metabolism have been intensively investigated over the past decades^1^. However, little is known about sulphur sensing and transducing signalling networks in plants, mainly hindered by the innate capacity of plant cells to produce myriad complexity of sulphur-containing amino acids and compounds via sulphur assimilation and metabolic pathways.

The evolutionarily conserved target of rapamycin (TOR) kinase acts as a master regulator that coordinates cell proliferation and growth by integrating nutrient, energy, hormone and stress signals in all eukaryotes. In plants, TOR functions as a central hub that integrates various nutrient signals, e.g. carbon, nitrogen, amino acids and phosphorus availability to coordinate growth and development^4-8^. Recently, the relationship between TOR and sulphur signalling has also emerged^9^. Sulphur assimilation begins with sulphate that is absorbed by sulphate transporters in the roots, and then transformed into adenosine 5’-phosphosulphate (APS), sulphite, and sulphide, which are catalysed by ATP sulfurylase (ATPS), APS reductase (APR), and sulphite reductase (SIR) respectively^3^. In the *sir1-1* mutant, which could hardly produce sulphide, TOR activity is down-regulated, and glucose level is significantly lower than that in wild-type *Arabidopsis*^10^. Interestingly, exogenous supply of glucose or reducing glutathione synthesis by inhibiting the glutamate-cysteine ligase activity partially restores the dwarf growth phenotype and increase TOR activity in the *sir1-1* mutant^10,11^, suggesting that sulphur limitation caused by decreasing of sulphite reduction to sulphide might pinpoint on glucose energy signalling to affect TOR activity. However, the connection between the direct sulphate and TOR signalling is not explored yet.

In plant development, cell proliferation in the leaf primordium of shoot apices is critical for the production of new leaves, and is strictly controlled by internal developmental cues, nutrients, hormones, and external environmental signals^12,13^. We previously showed that TOR can sense and coordinate energy (glucose), hormone (auxin) and environmental (light) signals in leaf primordium to activate cell proliferation and true leaf development^4,13^. In this study, we discovered that sulphate starvation strongly inhibited TOR activity and abolished cell proliferation in the leaf primordium even in the presence of exogenously supplied glucose. Interestingly, this low TOR activity could be rescued by low concentration of inorganic sulphate but not organic cysteine (Cys) or glutathione (GSH), indicating that sulphate could function as a direct nutrient signal to activate TOR and stimulate cell proliferation in the leaf primordium for true leaf development. To better understand the molecular landscape of this novel sulphate-TOR signalling network in shoot apex activation, we performed RNA-seq analyses to identify the sulphate-TOR directed transcriptome in the shoot apex. This tissue specific transcriptome analyses uncovered previously unknown sulphate-orchestrating genes and demonstrated that, as the central growth integrator, TOR kinase can sense different upstream nutrient signals to control both shared and differential transcriptome networks for precisely modulating plant proliferation, growth and stress responses.

## Results

### Sulphate availability activates TOR in a glucose-energy independent pathway

It was reported that plastidic sulphide level could indirectly influence TOR activity by interrupting the glucose-energy signalling since diminished sulphite reduction in *sir1-1* decreased the soluble sugar level and strongly impaired the TOR kinase activity and seedling growth, which could be recovered by exogenously resupplied glucose or sulphide^10^. However, sulphate deprivation strongly impaired plant growth and development, especially the shoot growth, e.g. the true leaf development, both in wild type (WT) and the *sir1-1* mutant, even in the presence of exogenously supplied glucose^14^ (Fig. 1a and b). Based on the well-established TOR kinase activity marker, phosphorylation level of T449 in the TOR conserved substrate ribosomal S6 kinase 1 (S6K1)^15^, we found that sulphate starvation dramatically reduced *Arabidopsis* TOR activity in the shoot apex, while the TOR mRNA or protein level was not affected (Fig. 1c-e). Furthermore, sulphate treatment effectively reactivated TOR both in the shoot apex of sulphate starved WT and *sir1-1* seedlings (Fig. 1f and g) and this activation was completely blocked by Torin2, a TOR specific inhibitor^13^, while the exogenously treatment of glucose, even at the high concentration (3%), still cannot reactivate TOR activity in sulphate starved seedlings (Fig. 1h). These results suggested that there is another sulphate-activated TOR pathway, which is independent from glucose availability.

**Figure 1.**
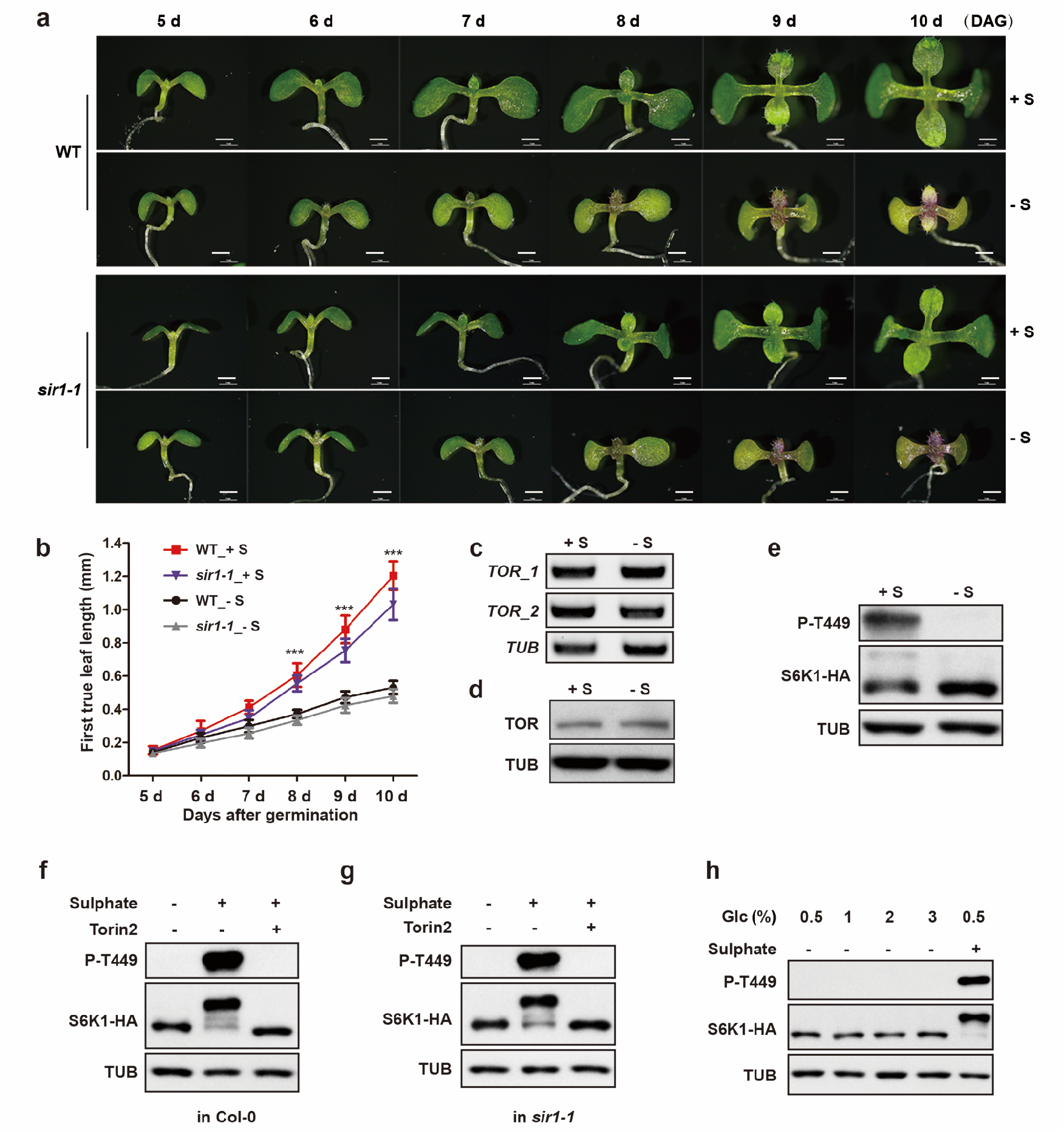
Sulphate availability regulates shoot apex development and TOR activity in a glucose-energy independent pathway. **a** Sulphate starvation inhibited *Arabidopsis* true leaf development. DAG: days after germination. WT or *sir1-1* seedlings were geminated in 1/2 MS liquid medium with or without sulphate (S) for 5-10 d. Scale bar, 1 mm. **b** Quantification of the first true leaf size of (**a**). Data are mean ± SD; n > 20, unpaired two-tailed *t*-test; \*\*\**P* < 0.001. **c** TOR mRNA level in shoot apexes was not affected by sulphate starvation. Semi-quantitative RT-PCR analyses. Total RNA was isolated from shoot apexes. *TOR_1* and *TOR_2* indicate two pairs of primers specific for the *TOR* gene. **d** TOR protein level in shoot apexes was not affected by sulphate starvation. TOR protein level was detected by western blotting using TOR specific antibody, and anti-Tubulin (TUB) was used a loading control. **e** TOR activity in shoot apexes was significantly decreased in response to sulphate starvation. TOR kinase activity was analysed based on p-T449 of S6K1 by western blotting. Anti-Tubulin (TUB) and anti-HA (S6K1-HA) were used as loading controls. **f-g** Sulphate re-activated TOR in shoot apexes both in WT and the *sir1-1* mutant. The 7 d sulphate starved *35S::S6K1-HA* (**f**) or *35S::S6K1-HA/sir1-1* (**g**) seedlings were recovered by 1 mM sulphate for 1 h with or without pre-treated TOR inhibitor Torin2 (25 µM) for 1 h. **h** Glucose cannot replace sulphate to reactivate TOR. The 7 d sulphate starved *35S::S6K1-HA* seedlings were treated with 1%, 2% or 3% glucose for 1 h, 1 mM sulphate treatment as a positive control.

### Sulphate as a primary upstream signal to activate TOR

After entering plants, the inorganic sulphate is further converted into organic sulphur containing amino acids and compounds, such as cysteine (Cys) and glutathione (GSH) through sulphate assimilation pathway. We characterized whether these sulphur nutrients can activate TOR in shoot apexes of sulphate starved seedlings within different concentration or kinetics patterns. Interestingly, 10 µM sulphate (standard MS medium contains about 1500 µM sulphate) was already able to activate TOR efficiently (Fig. 2a and b). The time-course experiment showed that sulphate could trigger a significant activation of TOR at 30 min, and the activation reached its peak at 60 min (Fig. 2c and d). However, comparing to sulphate, Cys and GSH were less efficient, and higher concentration (100 µM) were need for fully TOR activation in shoot apexes of sulphate-starved seedlings (Fig. 2a-d). Moreover, another sulphur-containing amino acid methionine (Met), and glutamine (Gln) which was reported to activate TOR in nitrogen signal pathway^7^, cannot activate TOR even at high concentration (1 mM) (Fig. 2e).

**Figure 2.**
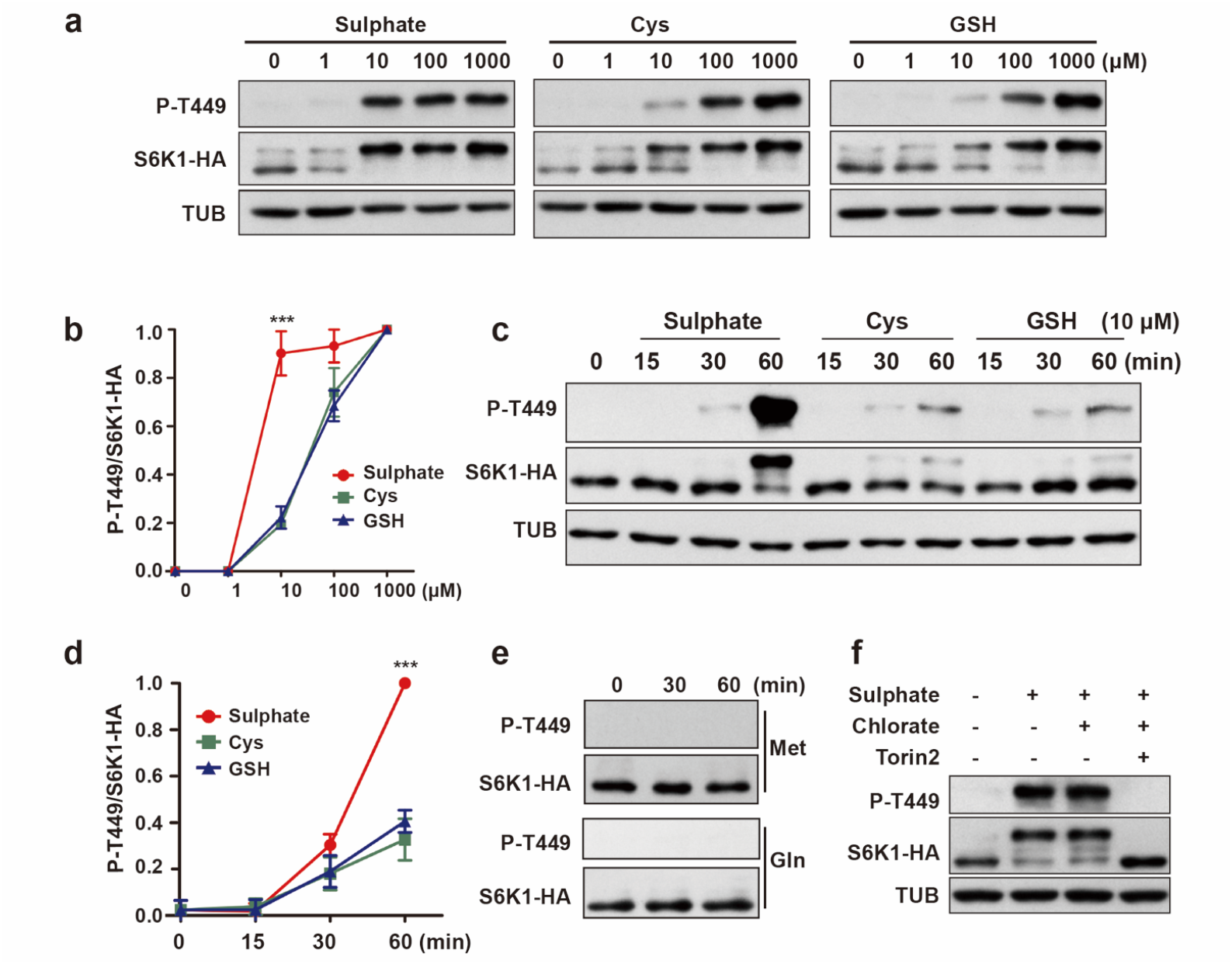
Sulphate functions as a primary upstream signal to activate TOR. **a** Concentration-dependent activation of TOR by different sulphur sources. *35S::S6K1-HA* seedlings were grown in sulphate-free medium for 7 d, then recovered by sulphate, cysteine (Cys) or glutathione (GSH) with the indicated concentration for 1 h. **b** Quantification of relative P-T449 intensity of (**a**), mean ± SD, n = 3, unpaired two-tailed *t*-test; \*\*\**P* < 0.001. **c** Time-dependent activation of TOR by different sulphur sources. The 7 d sulphate starved *35S::S6K1-HA* seedlings were recovered with 10 µM of different sulphur sources for indicated times. **d** Quantification of relative P-T449 intensity of (**c**), mean ± SD, n = 3, unpaired two-tailed *t*-test; \*\*\**P* < 0.001. **e** Methionine (Met) or glutamine (Gln) could not reactivate TOR activity. Sulphate starved *35S::S6K1-HA* seedlings were recovered with 1 mM Met or Gln for 1 h. **f** Chlorate treatment did not affect sulphate-reactivated TOR. The 7 d sulphate starved *35S::S6K1-HA* seedlings were pre-treated with chlorate (300 µM) for 5 h or Torin2 (25 µM) for 1 h, then recovered with 10 µM sulphate for 1 h.

We further investigated the potential primary signal role of sulphate in TOR activation by chemical genetic approach. During the sulphur assimilation, sulphate is first converted into APS by ATP sulfurylase (ATPS), which could be specifically inhibited by chemical inhibitor chlorate^16,17^. Significantly, we found that under chlorate treatment, sulphate could still efficiently activate TOR in shoot apexes of sulphate starved seedlings (Fig. 2f). Together, these results indicated inorganic sulphate itself instead of downstream organic sulphur-contained compounds could act as a primary nutrient signal to activate TOR and cell proliferation in the shoot apex.

### Sulphate activates cell proliferation in the shoot apex depending on TOR

We next investigated the downstream molecular mechanism of how sulphate promotes the true leaf development. Considering that true leaves are originally derived from leaf primordium and the activity of cell division is essential for leaf organogenesis, we examined the mitotic activity in leaf primordium based on the expression of the mitotic reporter, *proCYCB1;1::GUS*^13^. During the time period from the fifth to the tenth day after germination, the expression of *proCYCB1;1::GUS* continued to increase in the sulphate-supplied medium (Fig. 3a and b). However, in the sulphate-free medium, the expression of *proCYCB1;1::GUS* decreased significantly, and the GUS signal completely disappeared on the tenth day after germination (Fig. 3a and b), indicating that sulphate starvation strongly diminished the cell division activity in the shoot apex region.

**Figure 3.**
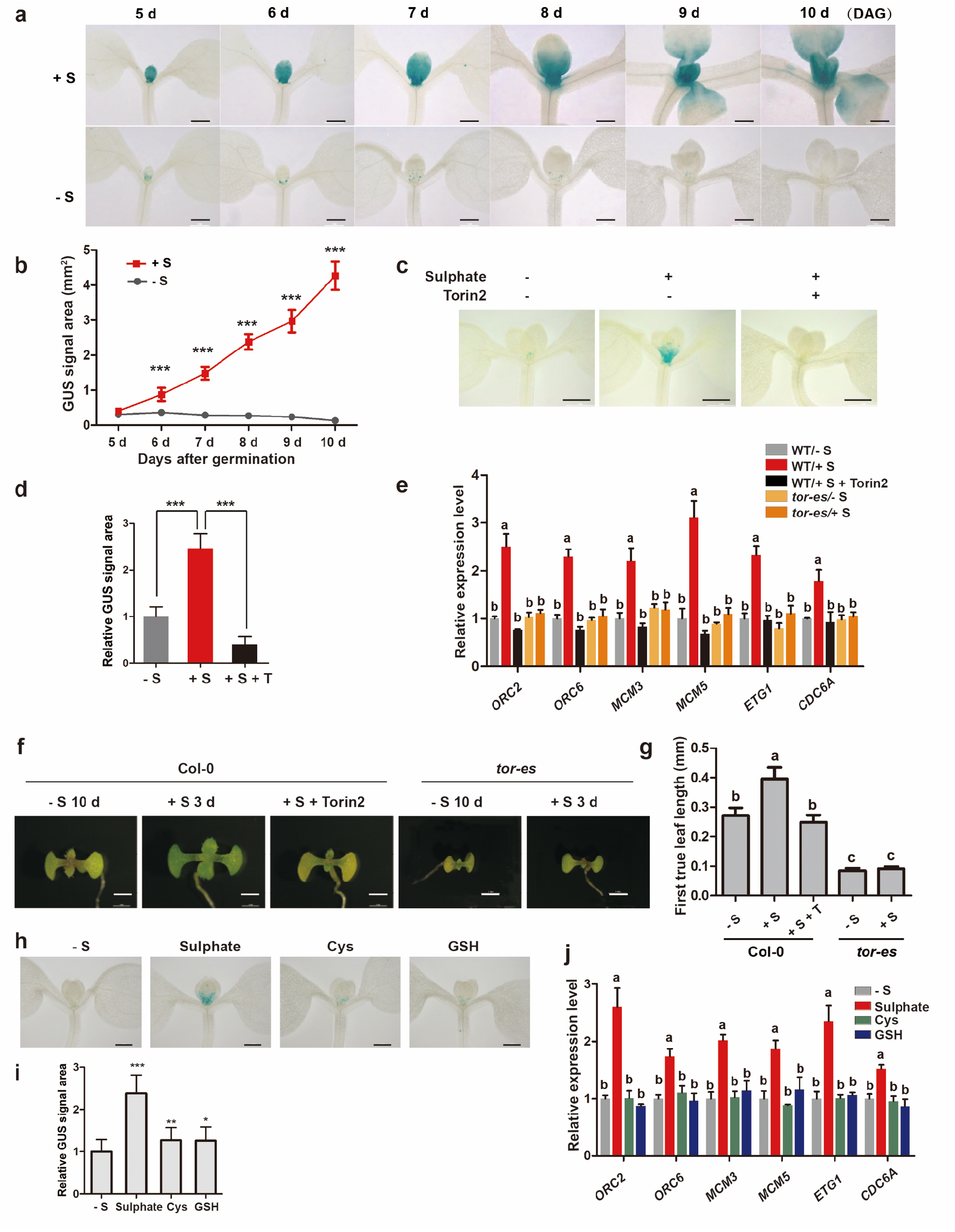
TOR is essential for sulphate-activated cell proliferation in shoot apex. **a** Sulphate starvation inhibited cell proliferation in *Arabidopsis* shoot apex indicated by GUS staining of *proCYCB1;1::GUS* seedling. *proCYCB1;1::GUS* seedlings were geminated in 1/2 MS liquid medium with or without sulphate (1 mM) for 5-10 d. DAG: days after germination. Scale bar, 1 mm. **b** Quantification of the GUS signal area of (**a**). Mean ± SD; n > 20, unpaired two-tailed *t*-test; \*\*\**P* < 0.001. **c** Inhibition of TOR blocked sulphate-reactivation of shoot apexes, indicated by GUS staining of *proCYCB1;1::GUS* seedlings. The 7 d sulphate starved *proCYCB1;1::GUS* seedlings were recovered by 1 mM sulphate for 9 h, with or without Torin2 (25 µM). Scale bar,1 mm. **d** Quantification of the GUS signal area of (**c**). Mean ± SD; n > 20, unpaired two-tailed *t*-test; \*\*\**P* < 0.001. **e** TOR inhibition blocked sulphate-reactivation of S-phase genes in shoot apexes. The 7 d sulphate starved WT or *tor-es* seedlings were treated with sulphate (1 mM) for 2 h, T: Torin2. QRT-PCR analyses. Means ± SD; n = 3, different letters represent significant difference between samples by one-way ANOVA (*P* < 0.05). **f** Sulphate-regulated reactivation of true leaf growth depending on TOR. Scale bar,1 mm. WT or *tor-es* seedlings were grown in sulphate-free medium for 7 d, then recovered with or without sulphate (1 mM) and Torin2 (25 µM) treatment for 3 d. **g** Quantification of the first true leaf size of (**f**). Mean ± SD; n > 20, different letters represent significant difference between samples by one-way ANOVA (*P* < 0.05). **h** Different potencies of sulphur sources for reactivation of cell proliferation in shoot apexes, indicated by GUS staining of *proCYCB1;1::GUS* seedlings. The 7 d sulphate starved *proCYCB1;1::GUS* seedlings were recovered by 10 µM of sulphate, Cys or GSH for 9 h. **i** Quantification of the GUS signal area of (**h)**. Mean ± SD; n > 20, unpaired two-tailed *t*-test; \*\*\**P* < 0.001, \*\**P* < 0.01, \**P* < 0.05. **j** Different potencies of sulphur sources for reactivation of S-phase genes in shoot apexes. QRT-PCR analyses. The 7 d sulphate starved WT seedlings were subjected to 10 µM of different sulphur sources for 2 h. Means ± SD; n = 3, different letters represent significant difference between samples by one-way ANOVA (*P* < 0.05).

To establish the direct functional link between sulphate nutrient and cell proliferation activity on shoot apex, we revealed that exogenously supplied sulphate (1 mM) can significantly reactivated the expression of *proCYCB1;1::GUS* in the sulphate starved seedlings (Fig. 3c and d). Since CYCB1;1 functions as a key regulator of G2/M phase transition within mitosis, we further examined the expression of S-phase cell cycle genes (*ORC2/6* (*ORIGIN RECOGNITION COMPLEX*), *MCM3/5* (*MINOCHROMOSOME MAINTENANCE*), *ETG1* (*E2F TARGET GENE*) and *CDC6A* (*CELL DIVISION CYCLE 6A*) in response to sulphate starvation. As shown in Fig. 3e, these S-phase cell cycle genes were also dramatically activated by 2 h sulphate treatment. Furthermore, sulphate-mediated reactivation of true leaf development, expression of *proCYCB1;1::GUS* and S-phase cell cycle genes were all strongly inhibited by Torin2 treatment or in the estradiol inducible *tor-es* mutant (Fig. 3c-g). In consistent with the different potency for TOR activation (Fig. 2a-d), the activation of *proCYCB1;1::GUS* and S-phase genes by sulphate was also stronger than that by Cys and GSH (Fig. 3h-j).

### Sulphate-TOR directed transcriptional network

To better understand the molecular landscape of the sulphate-TOR signalling network in the shoot apex activation, we performed RNA-seq analyses to compare transcriptome changes specifically in the shoot apex triggered by 2 h treatment of 10 µM sulphate, 10 µM Cys or 10 µM GSH in sulphate-starved seedlings with or without Torin2 (SRA BioProject ID: PRJNA678414) (Fig. 4a). Based on the relatively stringent statistics and filtering (*p* < 0.05, log_2_Fold change > 1 or < −1), we identified 754 up- and 1126 down-regulated genes differentially controlled by 10 µM sulphate in the *Arabidopsis* shoot apex (Fig. 4b and c, Supplemental Table 1), while only 14 and 6 genes were influenced by 10 µM GSH and 10 µM Cys, respectively (Fig. 4b and c, Supplemental Table 1). These specific and grand scope of gene expression changes indicated that sulphate orchestrates specific and significant transcriptional reprogramming. Strikingly, this swift global transcriptional reprogramming induced by sulphate was completely compromised in the presence of Torin2 treatment (Fig. 4c, Supplemental Table 1), supporting an indispensable role of TOR kinase in sulphate regulated transcriptome network.

**Figure 4.**
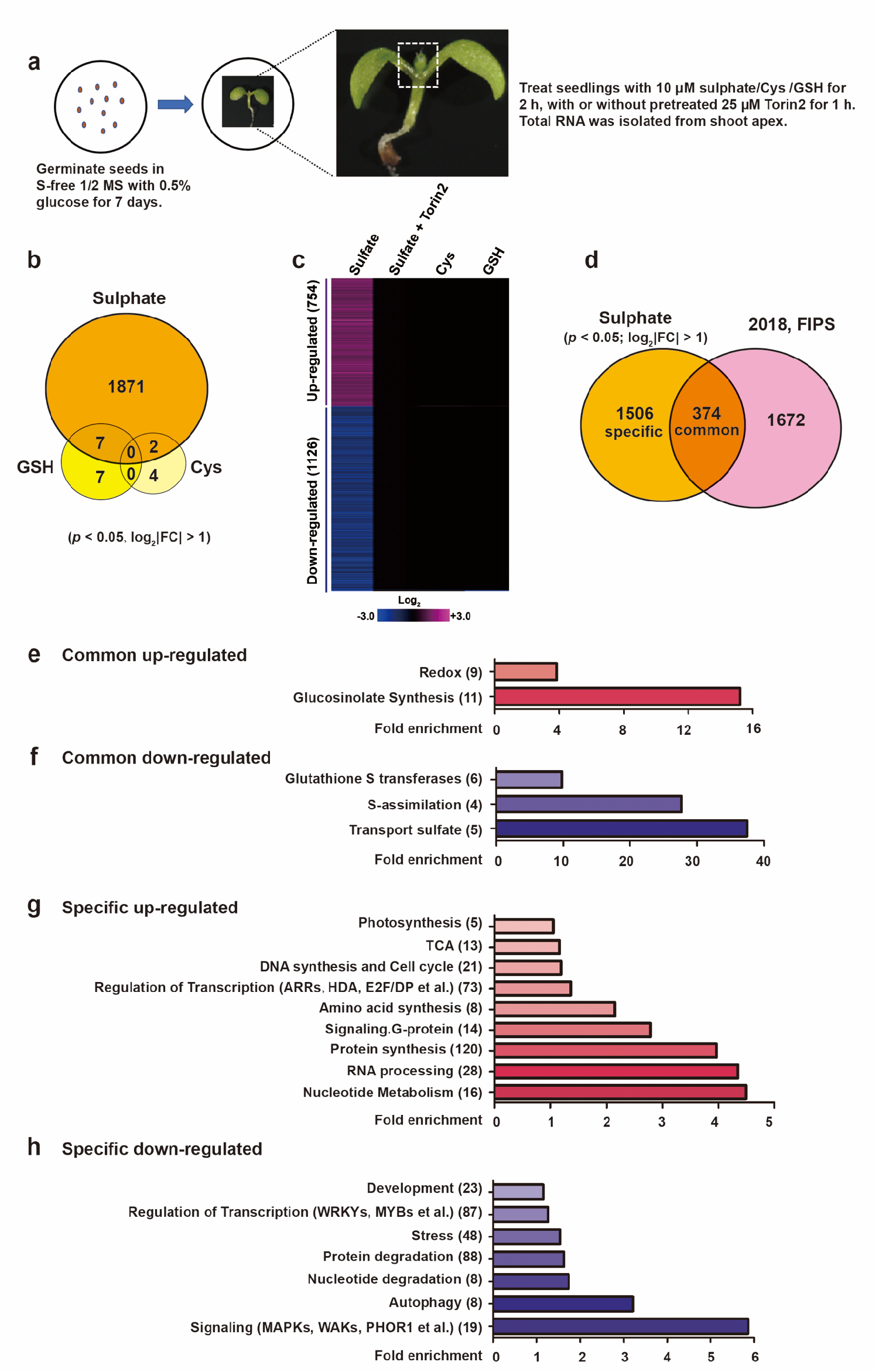
Sulphate-TOR orchestrates transcriptome reprogramming. **a** Schematic for preparing RNA-seq samples. **b** Comparison between sulphate, Cys and GSH specifically regulated transcriptomes (*p* < 0.05, log_2_|Fold change| >1). **c** Hierarchical clustering analysis of sulphate, sulphate with Torin2, Cys and GSH regulated genes. **d** Comparison between our new identified sulphate regulated genes with the classical sulphate-influenced genes. **e-f** Functional category analysis for common up- or down-regulated genes in (**d**). The numbers in brackets represent the number of genes in each category. **g-h** Functional category analysis for our new sulphate specific up- or down-regulated genes in (**d**). The numbers in brackets represent the number of genes in each category.

Recently, an integrative transcriptomic meta-analysis has been carried out based on five different sulphate starvation and recovering experiments, which were performed using a wide variety of sulphate concentrations, different growth systems (hydroponics or solid plates) and different tissues (roots and whole plants)^18^. There are total 2046 genes that significantly responded to the changes in sulphate availability in at least one experiment, and 27 core sulphate responding genes which were differentially expressed in all the five sulphate experiments^18^. We compared our RNA-seq data with those 2046 sulphur-response genes, and found that there were 374 common overlapped genes mainly involved in S-assimilation, redox, sulphur-related secondary metabolism and transport (Fig. 4d and e, Supplemental Table 2). Among the 27 core-sulphate responding genes identified by Henriquez et al, 2018, 25 genes were also presented in our RNA-seq analyses (Fig. S1), including genes responsible for sulphate transport and assimilation (*SULTR1;1, SULTR4;2* and *APR3*), and key repressors of GSLs metabolism (*SDI1/2, LSU-1, SHM7, GGCT2;1* [*ChaC-like*] and *BGLU28*) which negatively regulate the de novo biosynthesis of GSLs to save the available sulphur resources for primary metabolism^19,20^, reinforce that these genes are essential for sulphate response regardless of the growth stages and tissue/organ types. More strikingly, the unique experimental design and sensitivity of our system also facilitated the uncovering of 1506 previously unknown primary sulphate target genes especially enriched in activation of cell cycle and DNA synthesis, nucleotide metabolism, signalling, ribosomal biogenesis, protein synthesis and transcription, and repression of autophagy, nucleotide and protein degradation, and stress-related MAPKs, WAKs signalling and WRKY and MYB transcription activities (Fig. 4g and h, Fig. S1 and Supplemental Table 2).

### Comparison between sulphate-TOR and glucose-TOR directed transcriptional network

Glucose-TOR signalling has been reported to orchestrate global transcriptional reprogramming and promote root meristem activation and growth^4^. We further compared our sulphate-TOR and glucose-TOR regulated transcriptome (accession number: GSE114505). Although sulphate-TOR and glucose-TOR regulated transcriptomes were identified from different tissues (shoot apex and the whole 4 d seedlings, respectively), we found that 374 (50%) sulphate up-regulated genes and 210 (19%) sulphate down-regulated genes are also activated or repressed by glucose-TOR signalling, respectively (Fig. 5a and b). Remarkably, sulphate-TOR and glucose-TOR co-regulated genes stratify into specific regulatory and metabolic functional categories, including cell cycle and DNA synthesis, RNA transcription, ribosomal protein biogenesis, and protein synthesis machineries (Supplemental Table 3). For examples, BRIX superfamily members, which is essential for assembling large ribosomal subunits^21-23^, DExD-box helicases and transducing family protein requiring for pre-rRNA processing and modifications^24-26^, are all activated by both sulphate- and glucose-TOR signalling (Fig. 5c, Supplemental Table 3). These findings further strength a universal central role of TOR in controlling cell proliferation and protein translational processes.

**Figure 5.**
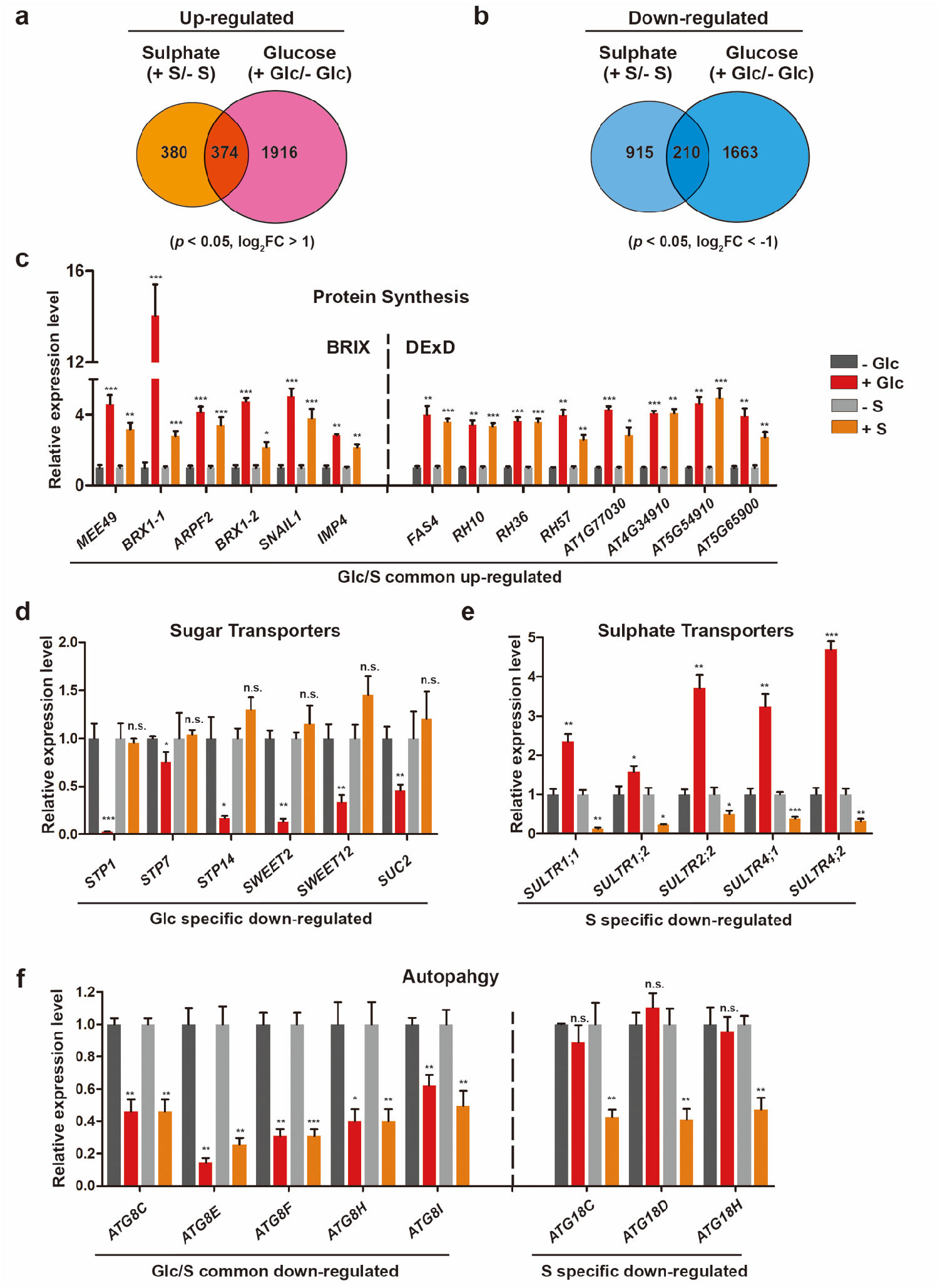
Comparison between sulphate-TOR and glucose-TOR directed transcriptional network. **a** Comparison between sulphate up-regulated genes with glucose up-regulated genes (*p* < 0.05, log_2_Fold change >1). **b** Comparison between sulphate down-regulated genes with glucose down-regulated genes (*p* < 0.05, log_2_Fold change < −1). **c-f** QRT-PCR analyses of different category genes. Means ± SD; n = 3; unpaired two-tailed *t*-test; \*\*\**P* < 0.001, \*\**P* < 0.01, \**P* < 0.05, n.s., no significance.

Notably, sulphate- and glucose-TOR signalling also orchestrate large sets of different gene categories. For example, unique to sulphate-TOR signalling was its pivotal roles in activating genes in sulphur-containing glucosinolates synthesis pathway, while glucose-TOR signalling also activated entire metabolic gene sets for lignin and flavonoid synthesis, and broad gene sets coding for the synthesis and modification of plant cell wall, cell wall proteins (arabinogalactan proteins and expansins), which were not identified in sulphate-TOR controlled transcriptome (Supplemental Table 3). The vast difference is also seen in the nutrient/metabolic transporters regulated by sulphate- and glucose-TOR signalling. The glucose-TOR signalling up-regulates genes involved in nitrate transport (*NRT1*.*1, NRT2*.*1, NRT2*.*3, NRT2*.*5, NRT2*.*6*), ammonium transport (*AMT1;2, AMT1;3, AMT1;4, AMT2;1*), phosphate transport (*ERD1, PHO1, PHT1;4, SHB1*) and sulphate transport (*SULTR2;2*) (Supplemental Table 3), but down-regulates genes for variously sugar transporters (*SUC2, STP1, STP7, STP14, SWEET2, SWEET12*) (Fig. 5d, Supplemental Table 3). By contrast, sulphate-TOR signalling specifically inhibits the expression of sulphate transporter (*SULTR1;1, SULTR1;2, SULTR2;2, SULTR4;1, SULTR4;2*) (Fig. 5e), which has been characterized as the classical sulphate starvation and recovering response genes^18,19^.

Autophagy is a process in which harmful or unwanted cellular components are delivered into lytic vacuoles to be recycled. The *ATG8s* and *ATG18s* gene family members are induced by various nutrient deprivation and environmental stresses, and play key roles in autophagy induction^5,27^. Sugar- or sulphur-starvation have been shown to trigger TOR-dependent autophagy for remobilization of internal nutrient resources^10,28^. Intriguingly, we found that sulphate-TOR negatively controls overlapped *ATG8s* genes (*ATG8C, E, F, H, I*) but unique *ATG18s* genes (*ATG18C, D, H*) compared to glucose-TOR signalling (Fig. 5f). Together, these findings uncover that TOR kinase can sense different upstream nutrient signals to control both shared and differential transcriptome networks for precisely modulating plant proliferation, growth and stress responses.

## Discussion

Sessile plants are challenged throughout their life cycles by various types of environmental stresses, such as nutrients deficiency, therefore, have to harmonize their complex metabolism, growth and developmental programs with the availability of nutrient. Sulphur is an essential macronutrient required for all eukaryotes, but how plants sense the sulphur nutrient level to adapt their growth remains largely unclear.

A recent study reported that sulphur limitation by decreasing sulphite reduction to sulphide in the *sir1-1* mutant affects the sugar content in plants, and then regulates plant growth and development through the TOR signalling pathway^10^. However, in this study, we found that sulphate deprivation strongly impaired *Arabidopsis* true leaf development, even with the high level of glucose supply (Fig. 1h). Sulphate treatment can effectively restore shoot growth and reactivate TOR kinase activity both in sulphate starved WT and *sir1-1* seedlings, and cannot be inhibited by the APS enzyme inhibitor chlorate (Fig. 1f and g, Fig. 2f). Strikingly, sulphate is more competent for activation of TOR activity, cell cycle gene expression and true leaf development, when compared to its downstream sulphur-contained metabolite GSH, and amino acids Cys and Met (Fig. 2a-e, Fig. 3h and i). Moreover, only a relatively small number of genes is overlap between sulphate-TOR- and glucose-TOR target genes (Fig. 5a and b). Therefore, our data strongly supports that sulphate might act as a primary nutrient signal to activate TOR, and is independent from glucose availability and downstream metabolic products of sulphur assimilation.

How sulphate activates TOR is still unclear and needs further research. In addition to sulphate, Cys and GSH can also activate TOR, although a higher concentration is required (Fig. 2a and b). Interestingly, 3-hydroxypropylglucosinolate, a defence-related glucosinolate, functions as a TOR inhibitor to block glucose-TOR promoted root meristem activation and root elongation^29^. These data indicate there are multifaceted functions of TOR intertwined with multiple layers of sulphur metabolic flux and its derived metabolites.

The evolutionarily conserved TOR kinase is a master nutrient sensor that coordinates cellular and organismal growth in all eukaryotes. In the past decades, especially in yeast and animals, TOR signalling has been linked mainly to amino acids and glucose nutrient sensing to modulate translational and transcriptional control^30,31^. However, different from animals who mainly obtain organic amino acids and carbon nutrients through dietary intake, sessile plants are autotrophic and uptake the essential inorganic macro- and micronutrients mainly from soil. In this study, we found that inorganic sulphate function as a primary upstream signal for TOR activation, which is uncoupled from sulphate assimilation pathway and glucose-mediated TOR activation. Couso et al. (2020) reported that in *Chlamydomonas*, phosphorus deprivation negatively affected LST8 protein stability, resulting in a down-regulation of TORC1 activity^8^. Therefore, plants might have evolved unique signalling mechanisms to sense and transduce these inorganic macro- or even micro-inorganic nutrients for TOR activation. Interestingly, in mammals and yeasts, TOR forms at least two structurally and functionally distinct protein complexes (TORCs) with both shared (LST8) and distinct (Raptor in TORC1, and Rictor, mSIN1 in TORC2), whereas only classical TORC1 is identified in plants^5^. In the future, it will be of great interest to explore whether plants possess functional TORC2, or even unique TORC to participate in plant specific biological processes, e.g. photosynthesis or inorganic nutrient sensing signalling networks.

## Materials and Methods

### Plant growth conditions

All *Arabidopsis* seeds were surface-sterilized by 75% ethanol with 0.05% Triton X-100 for 5 min and stored in sterile water for 2 d cold treatment in dark. The seeds were germinated in six-well plates containing 1 ml of liquid medium. Seedlings were grown in a plant growth chamber at 23°C light/21°C dark, 65% humidity, and 75 µmol/m^2^ s light intensity under 16-h light/8-h dark photoperiod.

### Plant materials

Col-0 was used as wild-type *Arabidopsis* plant. Mutant (*sir1-1*, GABI_550A09) and all transgenic plants are in Col-0 background. The estradiol-inducible *tor* RNAi line (*tor-es*), *35S::S6K1-HA* and *proCYCB1;1::GUS* were described previously^13,15^. The *35S::S6K1-HA* line was crossed with *sir1-1* line to generate *35S::S6K1-HA*/*sir1-1* line. Estradiol (10 µM) was used to induce TOR depletion, which was confirmed by a specific *Arabidopsis* TOR antibody^15^.

### Prepared sulphate starvation seedlings and sulphur recovery assay

To prepare the sulphate starvation seedlings, *Arabidopsis* seeds (7 seeds each well) were geminated in 6-well plates containing 1 ml of sulphate-free liquid medium (1/2 MS, pH = 5.7, Supplemental Table 4) for 7 d. The seedlings were then recovered by glucose, MgSO4, cysteine, glutathione, methionine or glutamine with the indicated concentration. The negative control (−) was treated with MgCl_2_ with the indicated concentration. For chemical treatments, sulphate starvation seedlings were pre-treated with Torin2 (25 µM) for 1 h or chlorate (300 µM) for 5 h before recovery experiments.

### Analyses of the shoot apex growth

For observing phenotype of the shoot apex growth, *Arabidopsis* seeds (7 seeds each well) were geminated in 6-well plates containing 1 ml of liquid medium (1/2 MS, pH = 5.7) with or without sulphate for 5-10 d. To analyse the TOR-dependent true leaf growth, *tor-es* seeds were planted in sulphur free liquid medium for 4 d, then treated with 10 µM estradiol for 3 d to induce *TOR* depletion^15^, following another 3 d sulphate recovery before phenotype observation.

All the photos were taken using a Leica stereoscopic microscope (Leica S8 APO). The representative shoot apexes were shown. The first true leaf length was measured with software Image J (NIH; https://imagej.nih.gov/ij/).

### GUS staining and quantification

The transgenic *proCYCB1;1::GUS* reporter line was used in this study^32^. Seven seedlings of each sample were treated with 1 ml staining solution (80 mM sodium phosphate buffer, pH = 7.0, 0.4 mM potassium ferricyanide, 0.4 mM potassium ferrocyanide, 8 mM EDTA, 0.05% Triton X-100, 0.8 mg/ml X-gluc (5-bromo-4-chloro-3-indolyl-β-D-glucuronide), 20% methanol) and incubated at 37°Cfor 12 h. Seedlings were distained by 75% ethanol at 37°Cfor 2 d, then visualized using Leica microscopy (Leica S9i). The representative seedlings were shown. The GUS staining area in the shoot apex was measured and quantified by the software Image J (NIH; https://imagej.nih.gov/ij/).

### Antibodies and western blot analysis

Phospho-p70 S6 kinase (p-Thr449) polyclonal antibody (Agrisera, AS132664) was used to detect TOR kinase phosphorylation of p-T449 in *Arabidopsis* S6K1. HA-tagged proteins were detected by anti-HA-HRP (sigma) monoclonal antibodies using standard techniques. Polyclonal *Arabidopsis* TOR antibody was generated as described^15^. The loading control Tubulin (TUB) was detected by monoclonal antibody anti-Tubulin (Abmart).

### Gene expression analyses

Total RNA was isolated from the shoot apex of seedlings with TRIzol (Invitrogen), except for the glucose starvation RNA samples which were isolated from whole 4 d glucose starved seedlings^4^. First-strand cDNA was synthesized from 0.5 µg total RNA with M-MLV Reverse Transcriptase (Promega). All qRT-PCR analyses were performed by CFX96 real-time PCR detection system with iQ SYBR green supermix (Bio-Rad). Semi-quantitative RT-PCR analyses was performed by PCR cycler (Bio-Rad). *TUB4* (*AT5G44340*) was used as control gene. All the primers used were listed in Supplemental Table 5.

### RNA-seq sample preparation, libraries construction and data analysis

The Col-0 seedlings were grown in sulphur free medium for 7 d, then were recovered by 10 µM sulphate, 10 µM Cys or 10 µM GSH for 2 h with or without pre-treated Torin2 (25 µM) for 1 h. Total RNA was isolated from shoot apex of seedlings.

The RNA-seq libraries were constructed with the NEB Next® Ultra™ II RNA Library Prep Kit for Illumina® following manufacturer’s instruction (NEB, Cat#E7775).

Libraries were sequenced on an Illumina Nova Seq 6000 (Paired-end 150 bp). RNA-seq data were then processed to identify differential expressed genes. Briefly, the sequence reads were aligned to *Arabidopsis* TAIR10 genome (release 48, ensemble plants, ftp://ftp.ensemblgenomes.org/pub/plants/release48/fasta/arabidopsis_thaliana/dna/) using Hisat2 (version 2.2.1) with default parameters^30^. After counting the gene expression with feature counts (version 2.0.1), the R package (version 4.0.0)-DESeq2 (version: 1.28.1) were introduced to normalize the reads by the median ratio method and calculate relative gene expression among different nutrient conditions with default parameters^33,34^. Moderated estimation of fold change and dispersion for RNA-seq data with DESeq2^34^. The significant differential expressed gene were identified with the parameter: log_2_|Fold change| >1, and adjusted *p* value (*padj*) < 0.05.

## Figures and figure legends

**Figure S1.**
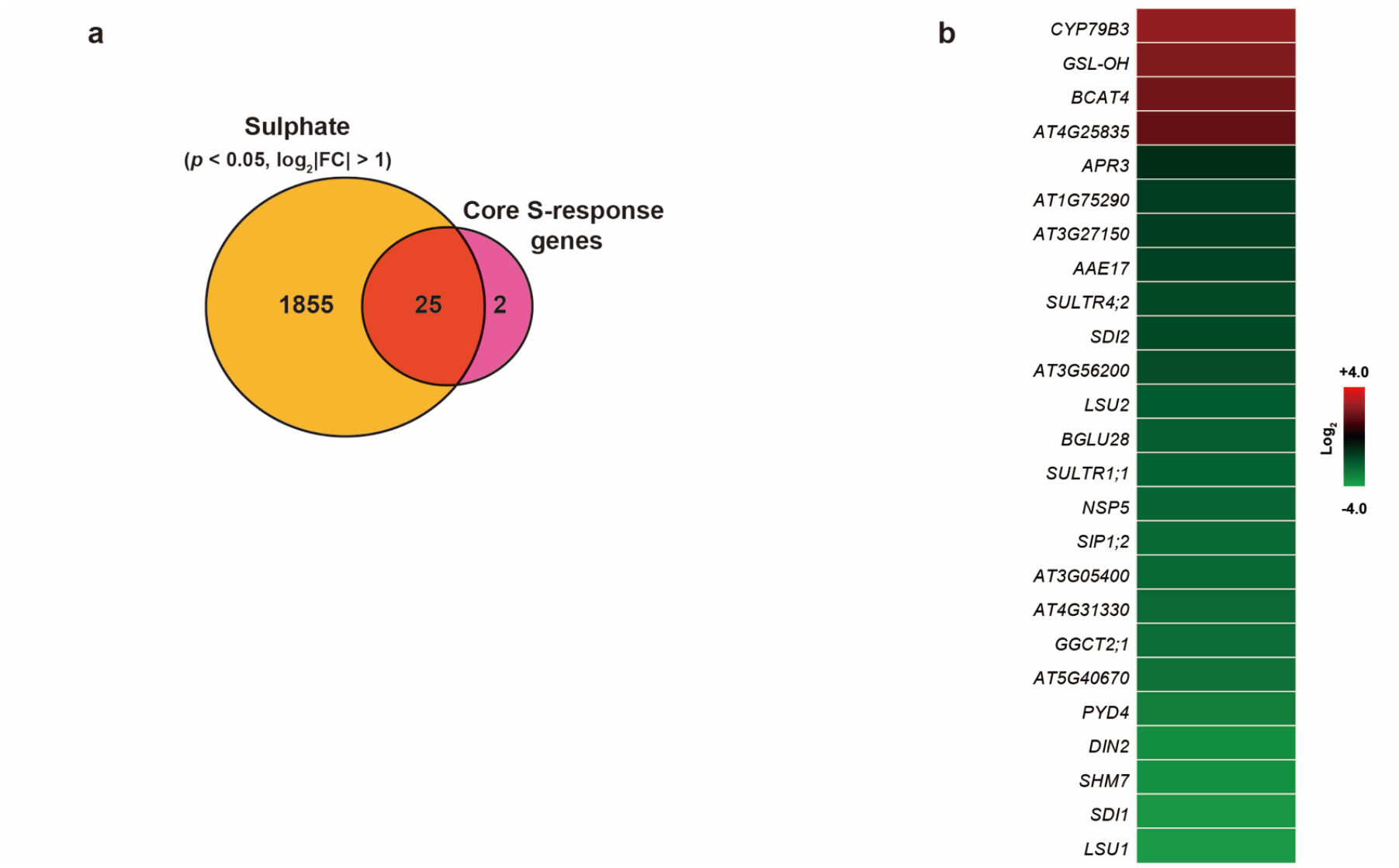
Sulphate regulates most of the core S-response genes in the shoot apex. **a** Comparison between sulphate regulated transcriptomes (*p* < 0.05, log_2_|Fold change| >1) and core S-response genes. **b** Heatmap of the 25 core S-response genes based on the RNA-seq data (Supplemental Table 1).

